# Record-matching of STR profiles with fragmentary genomic SNP data

**DOI:** 10.1101/2022.09.01.505545

**Authors:** Jaehee Kim, Noah A. Rosenberg

## Abstract

In many forensic settings, identity of a DNA sample is sought from poor-quality DNA, for which the typical STR loci tabulated in forensic databases are not possible to reliably genotype. Genome-wide SNPs, however, can potentially be genotyped from such samples via next-generation sequencing, so that queries can in principle compare SNP genotypes from DNA samples of interest to STR genotype profiles that represent proposed matches. We use genetic record-matching to evaluate the possibility of testing SNP profiles obtained from poor-quality DNA samples to identify exact and relatedness matches to STR profiles. Using 2,504 whole-genome sequences, we show that in some settings, similar match accuracies to those seen with full coverage of the genome are obtained by genetic record-matching for SNP data that represent 5-10% genomic coverage. Thus, if even a fraction of random genomic SNPs can be genotyped by next-generation sequencing, then the potential may exist to test the resulting genotype profiles for matches to profiles consisting exclusively of nonoverlapping STR loci. The result has implications in relation to criminal justice, mass disasters, missing-person cases, studies of ancient DNA, and genomic privacy.

## 1 Introduction

In a variety of contexts in forensic genetics, the identity of a DNA profile is sought from a biological sample with poor DNA quality, for which standard molecular techniques used with high-quality samples are unlikely to successfully produce genotypes. When the sample originates from trace sources such as burned, degraded, or ancient materials, only limited portions of the original genome might remain in the sample.

Routine genotyping of short-tandem-repeat loci (STRs) assumes that high-quality DNA samples contain DNA fragments in long sections of sequence. Hence, in a high-quality sample, the polymerase chain reaction can amplify the fragment that contains the entire section of DNA that lies between a specified pair of primer sequences [e.g., 1]. The amplification relies on the fact that both the primers and the fragment connecting them—which contains an STR region—are present in the DNA sample (Figure 1A).

**Figure 1:**
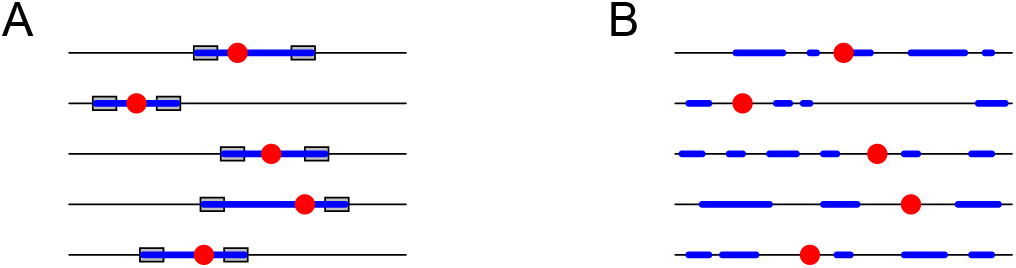
Genotyping of fragmented DNA might fail to amplify STRs, but it can amplify SNPs in the neighborhood of STRs. Each row depicts a chromosome, with an STR locus in red. The blue regions represent genotyped segments. (A) In high-quality DNA samples, STRs are genotyped by amplifying regions bracketed by PCR primers, depicted as gray boxes. (B) In low-quality DNA samples, PCR primers might not amplify, but some of the SNPs near an STR can be genotyped.

For degraded DNA samples, however, standard procedures for STR genotyping can be unlikely to succeed [2–4]. DNA fragments contained in the biological sample might be short and scattered, so that it is improbable that both primers and the full section of DNA that lies between them are present to be amplified. Nevertheless, although STR genotyping might fail, next-generation sequencing might be capable of producing genotypes of the available scattered DNA fragments (Figure 1B). Genetic information might be possible to extract from the sample, and in particular, genotypes might be possible to generate for some fraction of the single-nucleotide-polymorphism (SNP) sites in the genome [e.g., 5–12].

With next-generation sequencing of fragmentary materials, no particular genomic site can be reliably expected to appear in the genotype data. In particular, the STR loci that form the basis of standard forensic databases [13–15]—and that are genotyped by amplifying specific genomic sites—are unlikely to be obtained from the sample of interest, nor is any specific target set of SNPs. Thus, when an investigator seeks to query an unknown degraded sample for a genetic match to STR genotypes of a known individual or relative, or to search an STR profile database for a match, the fragmentary genotypes represent different and apparently incommensurable genetic loci from those available for potential matches.

Is it possible to identify genetic matches between a fragmentary SNP genotype profile obtained from a degraded DNA sample and the genotypes of a nonoverlapping set of STRs? In a technique termed “genetic record-matching,” we have recently demonstrated that, as a result of genotypic correlations that exist between STRs and their neighboring SNPs, it is frequently possible to identify matches between pairs of profiles, when one member of the pair is a SNP profile and the other is a forensic STR profile [16]. Furthermore, it can often be determined that a pair of profiles, one containing genome-wide SNPs and the other containing forensic STRs, represents a pair of close relatives [17].

In our calculations, however, we have made use of genome-wide SNP datasets with high genotyping quality, with a high level of genomic coverage in the neighborhood of each forensic STR locus. What if the SNP data were instead fragmentary, in the manner expected for degraded DNA and fragmented genotyping? This problem of record-matching between STR profiles and fragmentary SNP profiles represents any of several possible scenarios: matching the SNP profile of a degraded crime-scene sample to the STR profile of a specific known suspect, querying a degraded crime-scene SNP profile against a database of STR profiles, matching the SNP profile of an ancient DNA sample to specific STR profiles of possible living relatives, matching the SNP profile of a degraded DNA sample in a missing-persons or mass-disasters case to STR genotypes from known missing persons or their relatives, or querying it against an STR database of many potential candidates.

Here, we consider genetic record-matching between STRs and fragmentary SNP data, assuming that the available SNP data represent limited genomic coverage. We assess coverage levels from complete genomes down to 0.4% genomic coverage, evaluating the potential of record-matching for each level. We consider scenarios in which the hypothesis is that a SNP profile and an STR profile originate from the same person, from a parent–offspring pair, or from siblings. We find that for SNP and STR profiles from the same individual, the accuracy of genetic record-matching is often comparable with reduced coverage to that seen with the full genome. The results suggest a statistical approach for determining the identity of degraded DNA samples in criminal-justice, mass-disasters, missing-persons, and ancient-DNA scenarios, even without the potential for genotyping standard forensic STRs from those samples.

## 2 Materials and methods

### 2.1 Dataset

Because our method relies on the statistical connections between SNP and STR genotypes, we examine two well-studied repositories that contain genotype data of individuals typed at both SNP and STR loci. First, we use data from the Human Genome Diversity Panel (HGDP), as studied by [16] and [17]. This dataset contains unphased genotypes at 642,563 SNPs and 17 Codis STRs in 872 individuals from 52 worldwide populations. The Codis STRs are the 13 original Codis core loci and 4 of the 7 loci in the expanded set (D2S441, D10S1248, D19S433, and D22S1045).

The second dataset is a phased reference SNP–STR haplotype panel of Saini et al. [18] from the 1000 Genomes Project phase 3 [19, 20] with high-quality SNP genotypes obtained from whole-genome sequencing. The 1000 Genomes dataset contains 2,504 individuals from 26 worldwide populations typed at 11 of the 13 original Codis core loci—CSF1PO, D13S317, D18S51, D3S1358, D5S818, D7S820, D8S1179, FGA, TH01, TPOX, and vWA—and all 7 expanded Codis core loci [15], and genomic data at 27,185,239 SNPs.

Tables 1 and 2 compare the HGDP and 1000 Genomes datasets. The 1000 Genomes has higher SNP density in the neighborhood of each Codis STR than does the HGDP, with an average of ∼ 11,000 SNPs in a 1-Mb window centered at an STR locus in the 1000 Genomes compared to *∽*275 SNPs for HGDP.

**Table 1:**
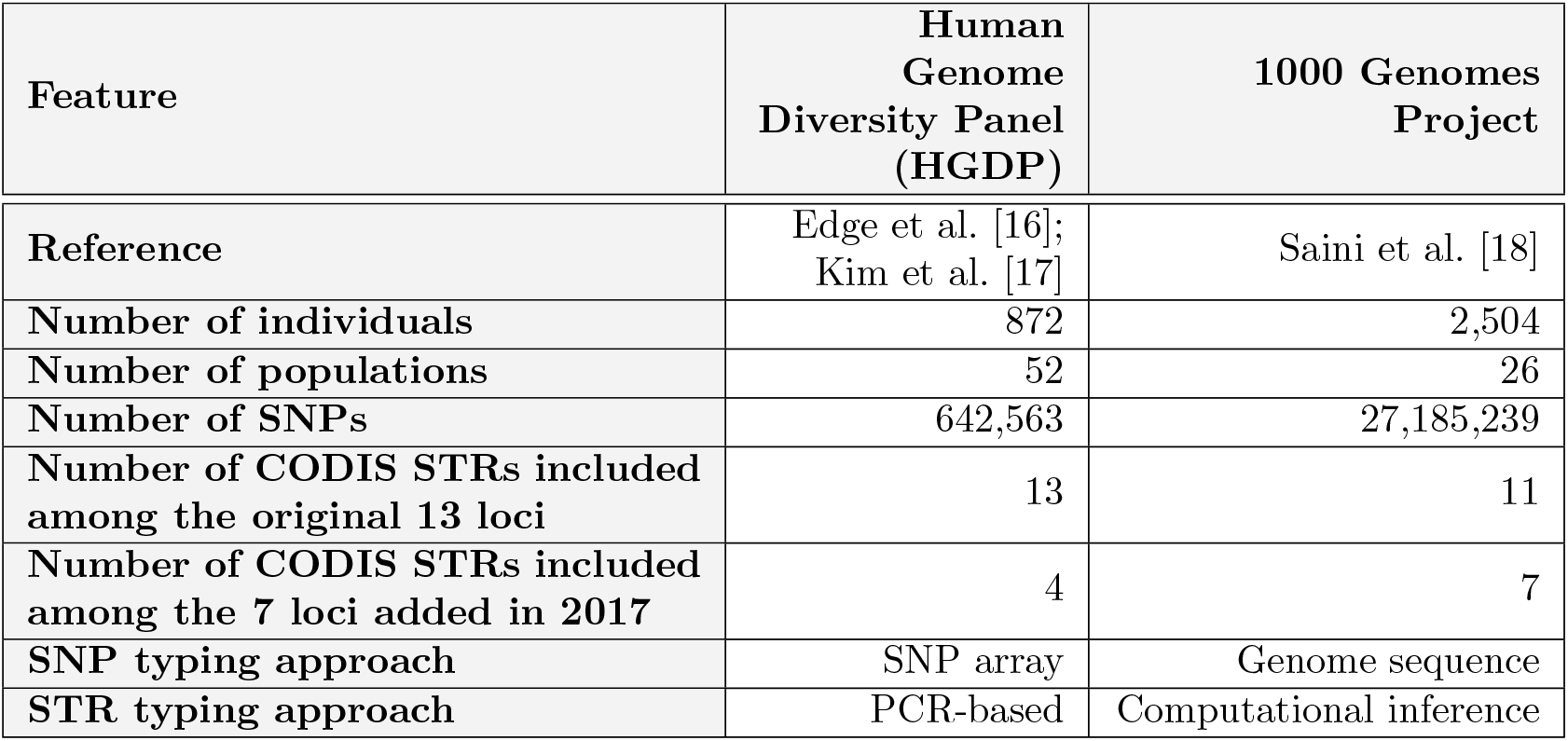
Summary of the features of two datasets used for joint analysis of CODIS STRs and genomic SNPs.

| Feature | Human Genome Diversity Panel (HGDP) | 1000 Genomes Project |
| --- | --- | --- |
| Reference | Edge et al. [16];<br>Kim et al. [17] | Saini et al. [18] |
| Number of individuals | 872 | 2,504 |
| Number of populations | 52 | 26 |
| Number of SNPs | 642,563 | 27,185,239 |
| Number of CODIS STRs included among the original 13 loci | 13 | 11 |
| Number of CODIS STRs included among the 7 loci added in 2017 | 4 | 7 |
| SNP typing approach | SNP array | Genome sequence |
| STR typing approach | PCR-based | Computational inference |

**Table 2:**
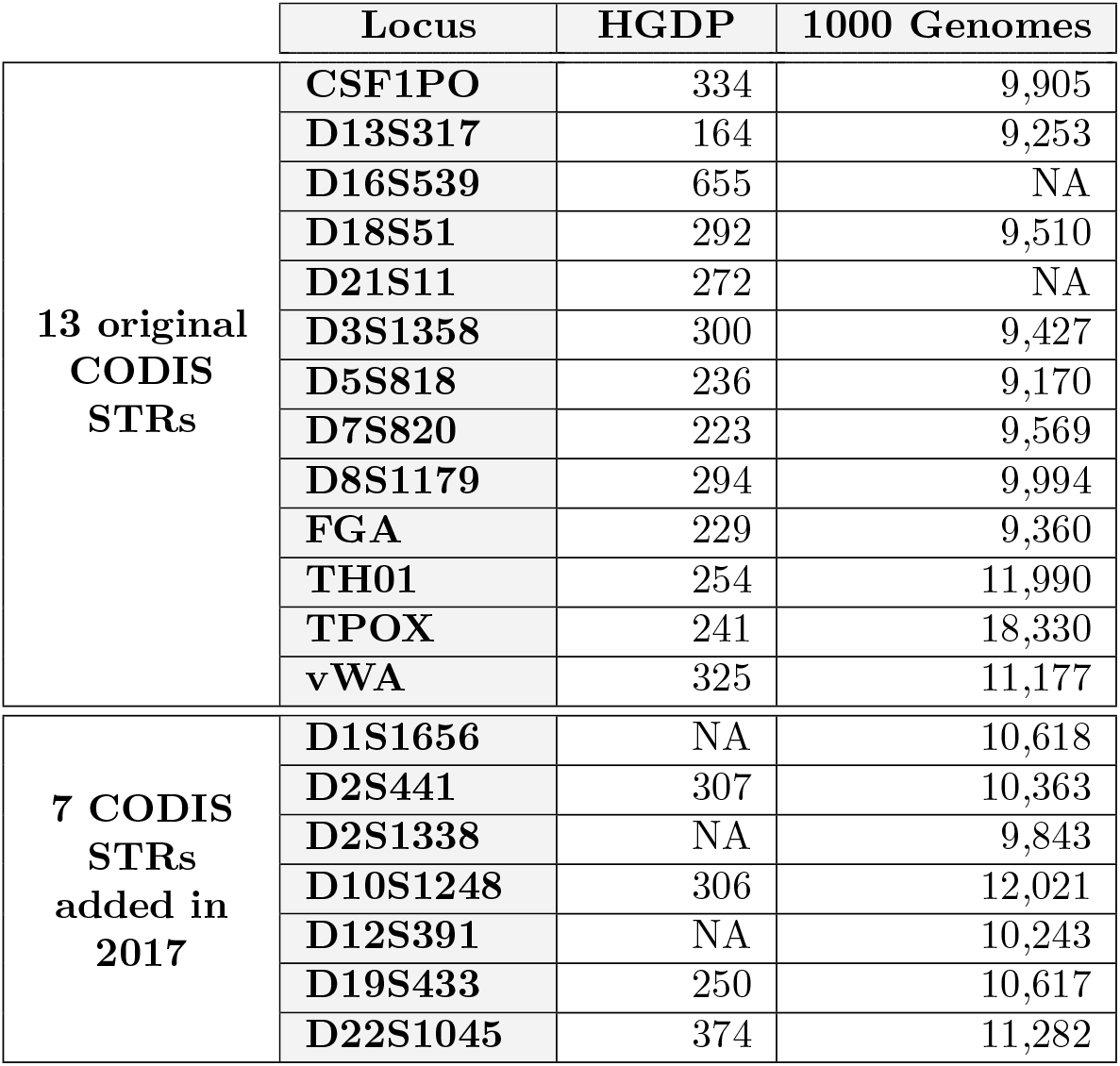
Number of SNPs in 1-Mb windows centered at the CODIS loci in the HGDP and 1000 Genomes datasets. The 15 STR loci used in our simulations are those loci for which data were present in both the HGDP and 1000 Genomes datasets.

|  | Locus | HGDP | 1000 Genomes |
| --- | --- | --- | --- |
| 13 original<br>CODIS<br>STRs | CSF1PO | 334 | 9,905 |
|  | D13S317 | 164 | 9,253 |
|  | D16S539 | 655 | NA |
|  | D18S51 | 292 | 9,510 |
|  | D21S11 | 272 | NA |
|  | D3S1358 | 300 | 9,427 |
|  | D5S818 | 236 | 9,170 |
|  | D7S820 | 223 | 9,569 |
|  | D8S1179 | 294 | 9,994 |
|  | FGA | 229 | 9,360 |
|  | TH01 | 254 | 11,990 |
|  | TPOX | 241 | 18,330 |
|  | vWA | 325 | 11,177 |
| 7 CODIS<br>STRs<br>added in<br>2017 | D1S1656 | NA | 10,618 |
|  | D2S441 | 307 | 10,363 |
|  | D2S1338 | NA | 9,843 |
|  | D10S1248 | 306 | 12,021 |
|  | D12S391 | NA | 10,243 |
|  | D19S433 | 250 | 10,617 |
|  | D22S1045 | 374 | 11,282 |

Our previous studies have analyzed familial record matching in the HGDP dataset [16, 17]. With more SNPs and individuals than the HGDP dataset, Saini et al. [18] demonstrated that the genotype imputation accuracies at Codis STRs from neighboring SNPs are slightly higher when using the 1000 Genomes (see their Table S2). We expect that SNP–STR familial record matching performed on the 1000 Genomes will have higher match accuracies than the previous observations with the HGDP. We further investigate the extent to which relatedness can be inferred between fragmentary SNP profiles and STR databases with the 1000 Genomes dataset as a reference panel from which the SNP–STR linkage information is drawn.

To enable comparisons of record-matching accuracies in the 1000 Genomes and HGDP datasets, we focus on the 15 Codis loci present in both datasets (Table 2)—11 from the original Codis STRs and 4 from the expanded Codis STRs—treating all individuals within a dataset as members of a shared population.

### 2.2 Genetic record matching

We examine familial relationship between a pair of individuals, one from an STR dataset and the other from a SNP dataset typed at specified genomic sequencing coverage.

#### 2.2.1 The relatedness match score

We follow [17] in computing match scores between profile pairs. In particular, for individual *i*, we represent the diploid genotype at STR locus *ℓ* by *R*_*il*_ and the diploid set of unphased genotypes at the neighboring SNP loci by *S*_*i*_. Considering *L* STR loci of individual *i*, we let *R*_*i*_ be the STR profile from the STR dataset, *R*_*i*_ = {*R*_*i*1_, *R*_*i*2_, …, *R*_*iL*_}, and we let *S*_*i*_ be the SNP profile from the SNP dataset, *S*_*i*_ = {*S*_*i*1_, *S*_*i*2_, …, *S*_*iL*_}.

In the absence of inbreeding, the familial relationship between two diploid individuals can be characterized by three coefficients, **Δ** = (Δ_0_, Δ_1_, Δ_2_), corresponding to the probabilities of three identity states *C*_0_, *C*_1_, *C*_2_ [21]. Each *C*_*k*_ represents a configuration in which, for the unordered diploid genotypes of two individuals at an autosomal locus, exactly *k* alleles are shared identically by descent between the individuals.

We test a specified relatedness hypothesis **Δ**_test_ between individual *A* with STR profile *R*_*A*_ and individual *B* with SNP profile *S*_*B*_ against a null model in which the individuals are unrelated. The test uses the log-likelihood-ratio relatedness match score comparing alternative and null hypotheses [17]:

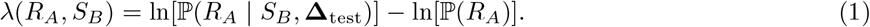

Assuming independence of the STR loci, Eq. 1 can be rewritten as:

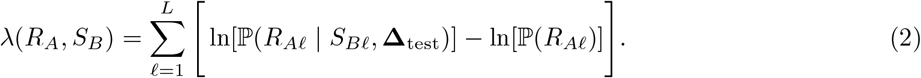

Likelihood ℙ (*R*_*Al*_ | *S*_*Bl*_, **Δ**_test_) can be decomposed over possible values of *R*_*Bl*_, the STR profile of individual *B* at locus *ℓ*:

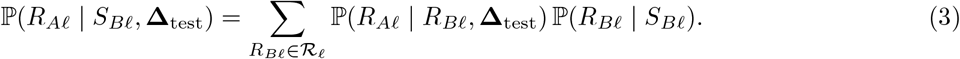

ℛ_*ℓ*_ denotes the set of possible genotypes at locus *ℓ*. ℙ (*R*_*Al*_ | *R*_*Bl*_, **Δ**_test_) is the probability of the observed STR genotype of individual *A* at locus *ℓ* conditional on a possible STR genotype of individual *B* at locus *ℓ* and the assumed relatedness hypothesis [21]. Evaluation of ℙ(*R*_*Al*_ | *R*_*Bl*_, **Δ**_test_) follows Kim et al. [17].

The ℙ (*R*_*Bl*_ | *S*_*Bl*_) term reflects the probability of possible STR genotypes of individual *B* at an STR locus *ℓ* conditional on the observed SNP profile surrounding the STR locus *ℓ* of individual *B*. We use `BEAGLE` and a phased SNP–STR haplotype reference to impute and obtain probabilities of unobserved genotypes at the STR locus *ℓ*, making use of the statistical association that exists between an STR and its neighboring SNPs. The detailed steps for this application of `BEAGLE` appear in Section 2.7.

Our record-matching pipeline can be applied to any familial relationships between *R*_*A*_ and *S*_*B*_, and we consider three such relationships: (1) same individual, **Δ**_true_ = (0, 0, 1); (2) parent–offspring, **Δ**_true_ = (0, 1, 0); and (3) sibling pairs, 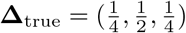.

#### 2.2.2 Prior and posterior odds

We report a number of results in terms of prior and posterior odds. Consider individual *A* with STR profile *R*_*A*_ and individual *B* with SNP profile *S*_*B*_, and two hypotheses,

*H*_0_ : *A* with STR profile *R*_*A*_ and *B* with SNP profile *S*_*B*_ are unrelated;

*H*_1_ : *A* with STR profile *R*_*A*_ and *B* with SNP profile *S*_*B*_ are related with relationship **Δ**.

Following [16], using Eq. 1, we can simplify the posterior odds for hypothesis *H*_1_:

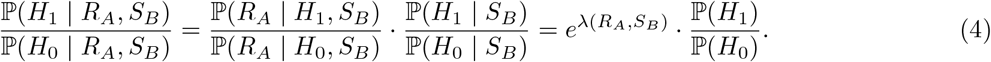

Here, *λ*(*R*_*A*_, *S*_*B*_) is a match score (Eq. 1) and ℙ (*H*_1_)/ ℙ (*H*_0_) is the prior odds for *H*_1_.

#### 2.2.3 Match assignment

We assigned matches from pairwise match scores as in [17]. For an STR dataset with *I*_*R*_ individuals and a SNP dataset with *I*_*S*_ individuals, we evaluated the relatedness match score under a test hypothesis **Δ**_test_ (Eq. 1) for all pairs of individuals, one with an STR profile and the other with a SNP profile. In our case, because STRs and SNPs are typed in the same individuals, we assume *I*_*R*_ = *I*_*S*_ = *I*.

We constructed a match-score matrix *M* of size *I* × *I*, where for all *j, k* in the set [*I*] = {1, 2, …, *I*}, *M*_*jk*_ = *λ*(*R*_*j*_, *S*_*k*_) is the entry for STR profile *R*_*j*_ from individual *j* and SNP profile *S*_*k*_ for individual *k*. From matrix *M*, we assigned matches by one of four schemes [16, 17]: one-to-one matching, one-to-many matching with a query SNP profile, one-to-many matching with a query STR profile, and needle-in-haystack matching.

In one-to-one matching, for a given row or a column of *M*, exactly one pair is selected as a match. We assigned matches via the Hungarian algorithm [22].

In one-to-many matching with a query profile (either STR or SNP), an observation in one dataset might be identified as having multiple relationship matches in the other dataset. When the query profile is an STR profile *R*_*j*_, its proposed match is *S*_*k*_, where *k* = argmax_*u∈*[*I*]_ *M*_*ju*_. When the query profile is a SNP profile *S*_*k*_, its proposed match is *R*_*j*_, where *j* = argmax_*u∈*[*I*]_ *M*_*uk*_.

Under both the one-to-one matching and one-to-many matching schemes, record-matching accuracy is defined as the fraction of pairs matched correctly among *I* true matches.

In needle-in-haystack matching, unlike in the other schemes, a database query is performed to locate a match only for one profile. In this setting, perfect accuracy is achieved when the match scores of all true matches exceed those of all non-matching profiles. The record-matching accuracy is defined as the proportion of true matches with greater match scores than the largest match score across all non-matching pairs.

### 2.3 Pedigree generation

To investigate familial record matching, following [17], we simulated random pedigrees from data on unrelated individuals. We drew mating pairs uniformly at random without replacement (ignoring population structure); each individual in a dataset appeared in exactly one mating pair. For each mating pair, we simulated two offspring. From each simulated pedigree, we randomly selected one of the parents and one of the offspring siblings for the parent–offspring scenario. We used both siblings for the sib-pair scenario.

In generating haplotypes of the offspring from parental haplotypes, following Kim et al. [17], we considered 1-Mb SNP windows extending 500 kb in each direction from a Codis locus midpoint, and we assumed that our 1-Mb window size was small enough that no recombination occurred within the windows. We treated Codis loci as independent, so that assortment was independent across Codis loci. Once pedigrees were generated, we randomized allele orders within individuals, discarding phase information, as the step in which we compute match scores in Section 2.2.1 begins with unphased data.

Note that prior to pedigree generation, for the HGDP dataset, which is unphased, we first used `BEAGLE` to phase the entire HGDP dataset to obtain individual haplotypes (Section 2.7). We then generated 10 random sets of 436 pedigrees from the 872 HGDP individuals following the above procedure.

The 1000 Genomes dataset contains phased haplotypes, so the initial phasing step prior to pedigree generation was omitted. Using 2,504 individuals in the 1000 Genomes dataset, we generated 10 random sets of pedigrees, each of which contains 1,252 simulated pedigrees.

### 2.4 Record matching with HGDP and 1000 Genomes datasets

We first evaluated HGDP and 1000 Genomes record-matching accuracies with the 15 Codis loci described in Section 2.1. Following Edge et al. [16] and Kim et al. [17], we split the data into disjoint training and test sets, with 75% of the individuals in the test set. We phased HGDP training sets using `BEAGLE` to obtain SNP–STR haplotypes that we used as a reference. Next, we again used `BEAGLE` for imputing STR genotypes from test-set SNP profiles with the phased SNP–STR haplotype reference panel from the training set (Section 2.7).

For the same-individual scenario, we generated 100 random partitions into training and test sets. When using the HGDP dataset, each partition contained 654 individuals in the training set and 218 in the test set. For 1000 Genomes, each partition contained 1,878 individuals in the training set and 626 in the test set. We estimated STR allele frequencies from the individuals in the training set.

For the parent–offspring scenario, for each of the 10 random sets of pedigrees, we generated 10 random partitions of the pedigrees into training and test sets, resulting in 100 random partitions of pedigrees into training and test sets in total. In each partition, the numbers of pedigrees in the training and test sets were 327 and 109, respectively, when using the HGDP dataset, and 939 and 313 for 1000 Genomes. For each test-set pedigree, without loss of generality, we assigned a SNP profile of a parent to the SNP dataset and an STR profile of an offspring to the STR dataset.

For the sib-pair scenario, we generated 100 random partitions of pedigrees into training and test sets—10 replicates for each of the 10 random pedigree sets. For each test-set pedigree, we randomly placed one sibling in the SNP dataset and the other in the STR dataset. In both parent–offspring and sib-pair scenarios, we estimated STR allele frequencies from the parents in training-set pedigrees.

For all three scenarios (same-individual, parent–offspring, sib-pair), for each of the 100 partitions, we constructed the match-score matrix from the test set and assessed record-matching accuracies for the four matching schemes (Section 2.2.3). The median, minimum, and maximum record-matching accuracies of the 100 replicates using the HGDP dataset are shown in Table S1. The corresponding values using the 1000 Genomes are shown in Table 3 when **Δ**_true_ = **Δ**_test_, and in Table S2 when **Δ**_true_ ≠ **Δ**_test_. The median value was chosen to be the lesser of two possible median values when the number of unique values was even.

**Table 3:** Record-matching accuracies using the 1000 Genomes dataset and 15 CODIS loci, for Δ_true_ = Δ_test_. The table summarizes 100 partitions into training and test sets, applying record matching to the 1000 Genomes dataset with no missing data. The STRs used are listed in Table 2.

| Same individual |  | Parent-offspring |  | Sib pairs |  | Match-assignment scheme |
| --- | --- | --- | --- | --- | --- | --- |
| Median | Min, Max | Median | Min, Max | Median | Min, Max |  |
| 1.000 | 1.000, 1.000 | 0.738 | 0.681, 0.812 | 0.693 | 0.623, 0.773 | One-to-one |
| 1.000 | 0.998, 1.000 | 0.649 | 0.581, 0.700 | 0.626 | 0.546, 0.674 | One-to-many: SNP query |
| 1.000 | 0.998, 1.000 | 0.649 | 0.597, 0.719 | 0.636 | 0.572, 0.703 | One-to-many: STR query |
| 0.992 | 0.941, 1.000 | 0.112 | 0.016, 0.256 | 0.160 | 0.019, 0.275 | Needle-in-haystack |

### 2.5 Simulation of fragmentary genomic SNP data

To generate random fragmentary genomic SNPs from the 1000 Genomes dataset, for each of the three relatedness scenarios, we selected a partition corresponding to the median one-to-one match accuracy with **Δ**_true_ = **Δ**_test_ (Section 2.4, Table 3). The match accuracy varies discretely across partitions, as it measures a fraction of test-data individuals correctly matched; when multiple partitions all produce the median-accuracy value, we picked one at random. Note that because the HGDP dataset contains far fewer SNPs than the 1000 Genomes dataset (∼2.5%), we conducted simulations of fragmentary SNP data for the 1000 Genomes only.

Under each choice of relatedness, from the full-coverage SNP profiles of individuals in the median-accuracy test set, we simulated fragmentary SNP data. For the same-individual scenario, this test set has 626 individuals; for the parent–offspring and the sib-pair scenarios, the test set has 313 individuals, one for each test-set pedigree as described in Section 2.4.

Among *N*_all_ = 27, 185, 239 SNPs in the 1000 Genomes dataset, the number in the 1-Mb windows around the *L* = 15 STR loci considered was *N*_win_ = 161, 968 (Table 2). We considered 30 values of genomic sequencing coverage *c*: {0.004, 0.006, 0.008, 0.01, 0.02, …, 0.19, 0.2, 0.3, …, 0.8, 0.9}. With the partition of the data into training and test sets fixed, for each *c*, we generated 100 random sets of fragmentary SNP data of the test-set individuals as follows. A schematic of the simulation pipeline appears in Figure 2.

**Figure 2:**
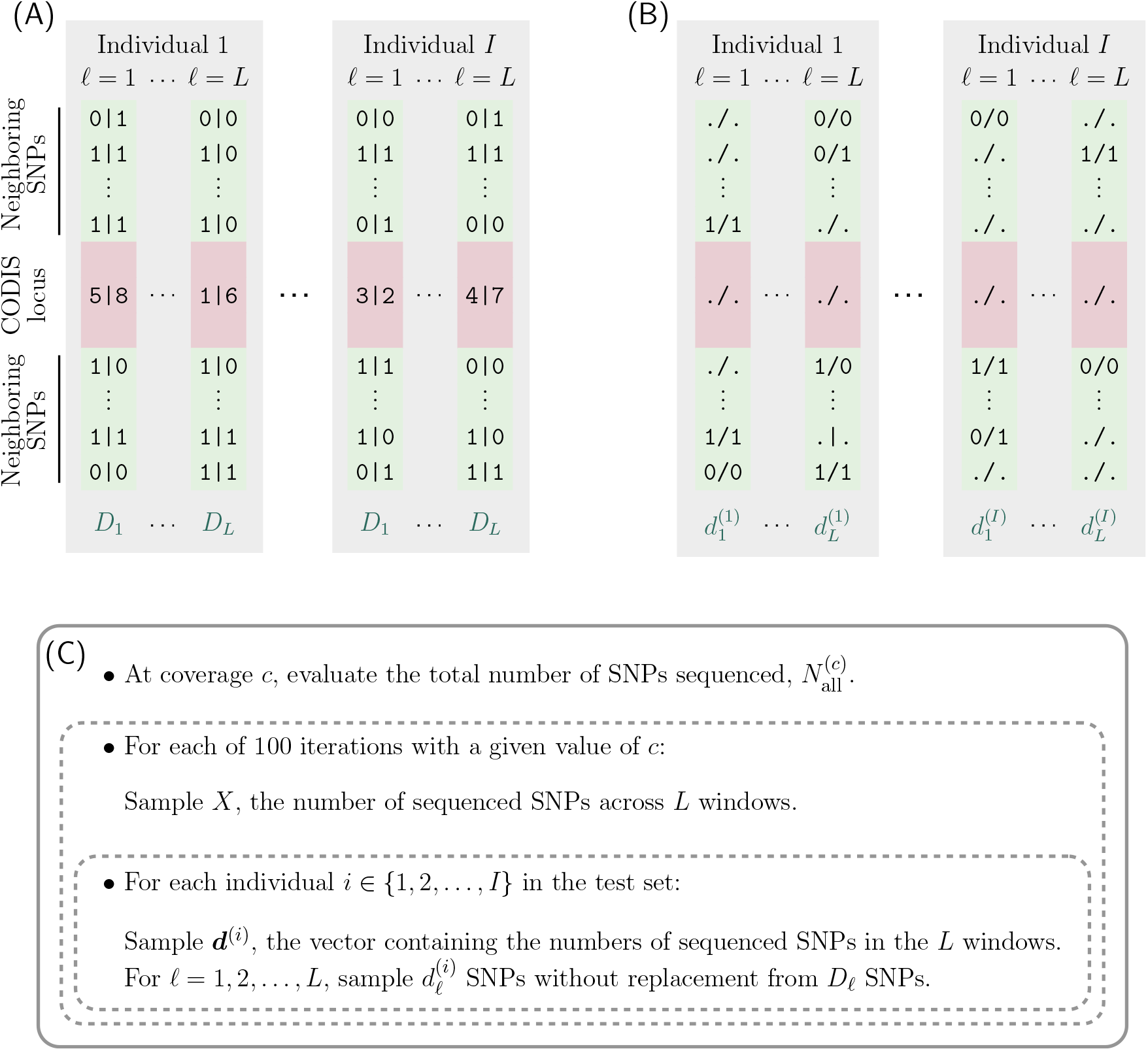
Schematic for simulating fragmentary SNP datasets for the individuals in the test set. (A) An example of the SNPs in a 1-Mb window (green) of a Codis locus (red) in two specific individuals. We denote the total number of SNPs in the whole genome with full coverage (*c* = 1) by *N*_all_ = 27, 185, 239. *D* (*ℓ* = 1, 2, …, *L*) indicates the number of SNPs in the 1-Mb window of the *ℓ*th Codis locus, and 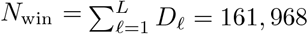, 968 represents the number of SNPs in all *L* 1-Mb windows (Table 2). (B) The simulated set of fragmentary SNPs for the individuals in (A). (C) The simulation pipeline for generating simulated fragmentary SNPs from the 1000 Genomes dataset. For a given sequencing coverage *c*, the total number of SNPs sequenced from the whole genome is 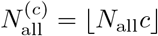. For a given sequencing coverage *c*, we repeat the following procedure 100 times to generate 100 random sets of fragmentary SNPs. We first sample *X*, the number of sequenced SNPs in *L* 1-Mb windows combined, from a binomial distribution with parameters 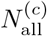 and *f* = *N*_win_/*N*_all_ ≈ 0.006. Using the sampled value of *X*, for each test individual *i* (*i* = 1, 2, …, *I*), we generate random sets of sequenced SNPs in the 1-Mb windows by first sampling individual-specific 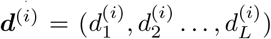—the vector of numbers of sequenced SNPs from each of the *L* windows—from a multinomial distribution with parameters *X* and (*D*_1_*/N*_win_, *D*_2_*/N*_win_, …, *D*_*L*_*/N*_win_). For each *ℓ* (*ℓ* = 1, 2, …, *L*), we then sample 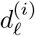 SNPs uniformly at random without replacement from the *D* SNPs of the full-coverage set.

We denote the number of SNPs within the 1-Mb window around the *ℓ*th Codis locus by *D*_*ℓ*_, with 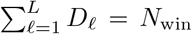, and we denote its relative proportion by *pl* = *D*_*ℓ*_/*N*_win_. The values of *D*_*ℓ*_ are listed in Table 2. Of *N*_all_ SNPs, the fraction of SNPs present in the *L* windows is *f* = *N*_win_/*N*_all_ ≈ 0.006.

For a given coverage *c*, the total number of SNPs sequenced is 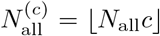. Assuming that all SNPs have equal probability of being sequenced, each sequenced SNP lies in one of the *L* 1-Mb windows around the Codis loci with probability *f*. For each simulated fragmentary SNP dataset with coverage *c*, we sampled *X*—a total number of SNPs sequenced in the *L* 1-Mb windows combined—from a binomial distribution with parameters 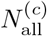 and *f*. For each test individual *i* in a simulated fragmentary dataset with *X* SNPs sequenced in the *L* windows combined, we sampled a vector 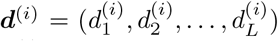 from a multinomial distribution with parameters *X* and ***p*** = (*p*_1_, *p*_2_, …, *p*_*L*_). Here, 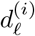 represents a number of SNPs sequenced within the 1-Mb window of the *ℓ*th Codis locus in a fragmentary SNP data of individual *i*. Once ***d***^(*i*)^ was sampled, for each 1-Mb window around the *ℓ*th Codis locus, we sampled 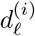 SNPs uniformly at random without replacement from *D_ℓ_* SNPs in the full-coverage dataset. As an example, Figure 2B displays a simulated fragmentary unphased genotype dataset for individuals in a test set shown in Figure 2A.

### 2.6 Record-matching of STR profiles with fragmentary genomic SNP data

Our record matching formulation introduced in Section 2.2 assumes no constraints on genomic coverage. Thus, we applied the record-matching pipeline to each simulated fragmentary SNP dataset of the test-set individuals.

For this analysis, as noted in Section 2.5, we fixed the training set at the median-accuracy partition generated in Section 2.4; it contained the full-coverage SNP–STR haplotypes of the training-set individuals. Hence, each relatedness scenario has a training set; we used this same shared training set across all 100 simulated fragmentary SNP datasets for that scenario.

We used the training set as a reference panel when imputing test-set STR profiles from their neighboring fragmentary SNP profiles according to Eq. 3. We also computed STR allele frequencies from the training set in order to compute the match score (Eq. 1).

For the same-individual scenario, the training set contained 1,878 individuals and the test set had 626. For each of 100 simulated fragmentary SNP datasets at a given genomic coverage *c*, we computed match scores of all pairs—one with a SNP profile and the other with an STR profile—and obtained a 626 × 626 match-score matrix. We then computed match accuracies under four matching schemes described in Section 2.2.3.

For the parent–offspring scenario, the training set contained 1,878 individuals: all parental pairs of the 939 pedigrees in the training set for the median-accuracy partition with **Δ**_true_ **= Δ**_test_ = parent–offspring.

The corresponding median-accuracy test set contained the remaining 313 pedigrees (Section 2.4). For each of the 100 simulated fragmentary SNP datasets at genomic coverage *c*, we computed a 313 *×* 313 match-score matrix to match the fragmentary SNP profiles of offspring to the STR profiles of parents in the test set.

Finally, for sib pairs, the training set contained 1,878 individuals, all parental pairs of the 939 pedigrees in the training set of the median-accuracy partition with **Δ**_true_ **= Δ**_test_ = sib pairs. The corresponding median-accuracy test set consisted of the remaining 313 pedigrees (Section 2.4). For each of the 100 simulated fragmentary SNP datasets at genomic coverage *c*, we computed a 313 *×* 313 match-score matrix to match fragmentary SNP profiles to sibling STR profiles in the test set.

### 2.7 BEAGLE settings for phasing

We used `BEAGLE` v. 5.0 [23, 24] for two purposes: phasing and imputation. First, in Section 2.3, we phased the entire HGDP dataset to generate the SNP–STR haplotypes from which we simulated random pedigrees (recall that this phasing step was not needed for the pre-phased 1000 Genomes dataset). In Section 2.4, we also phased unphased genotypes in HGDP training datasets.

For these analyses, we set `iterations=14`, `burnin=6`, `phase-states=280`, and `phase-segment=4.0`. We used `BEAGLE` default parameters shared between the phasing step and the imputation step in Section 2.8: `ne=1000000`, `err=0.0001`, `window=40.0`, `overlap=4.0`, `seed=-99999`, `step=0.1`, and `nsteps=7`.

### 2.8 BEAGLE settings for imputation

We also used `BEAGLE` v. 5.0 for estimating the unobserved STR genotype probabilities, P(*R*_*B*_ | *S*_*B*_) in Eq. 3, as in Section 2.2.1. This analysis, which we employed in Sections 2.4 and 2.6, used the phased training-set SNP–STR haplotypes as a reference and augmented them with the SNP genotypes of the unphased test set. We then estimated the genotype probabilities at the STR loci in the test set based on the neighboring SNPs.

This analysis used identical `BEAGLE` settings to those used in Section 2.7, with some exceptions. First, this analysis used `gp=true`, `impute=true`, `imp-states=1600`, `imp-segment=6.0`, `cluster=0.005`, and `ap=false`. Second, the analysis employed the human reference genome GRCh37 genetic map, whereas the phasing analysis did not use a genetic map.

## 3 Results

We first focus on correctly specified hypotheses, **Δ**_true_ **= Δ**_test_. Under three relatedness scenarios—same individual, parent–offspring, and sib-pair—we examine the case of **Δ**_true_ **= Δ**_test_ as a function of the SNP coverage *c*. This analysis follows the procedures in Section 2.6. Numerical summaries appear in Table 3.

### 3.1 Same individual

Figure 3A-D shows the record-matching accuracy for **Δ**_true_ **= Δ**_test_ = same individual. For each of four matching schemes, the figure shows the median accuracy for HGDP across partitions, producing slightly greater values than those obtained in corresponding analyses in Table 2 of our previous study [17], which used 13 rather than 15 loci. It also shows the median accuracy for the larger and denser 1000 Genomes dataset, and for 1000 Genomes datasets with fragmentary SNP data.

**Figure 3:**
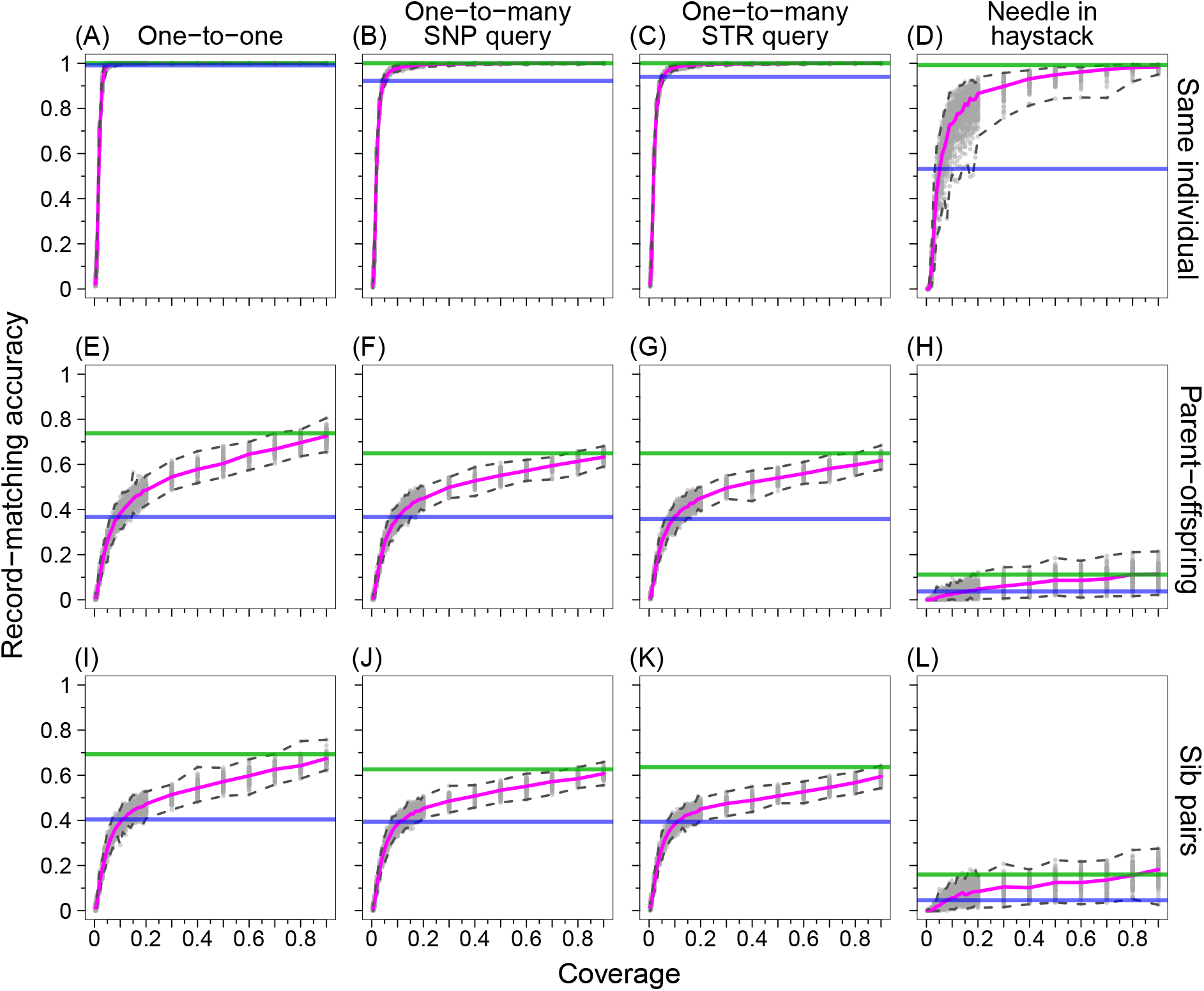
Record-matching accuracy in fragmented genomic data as a fraction of the genomic coverage *c*, for Δ_true_ = Δ_test_. **Δ**_true_ is the true relationship between pairs of individuals, and **Δ**_test_ is the test relationship hypothesis on the basis of which match scores are computed. (A) Same individual, one-to-one matching. (B) Same individual, one-to-many matching with a query SNP profile. (C) Same individual, one-to-many matching with a query STR profile. (D) Same individual, needle-in-haystack matching. (E) Parent– offspring, one-to-one matching. (F) Parent–offspring, one-to-many matching with a query SNP profile. (G) Parent–offspring, one-to-many matching with a query STR profile. (H) Parent–offspring, needle-in-haystack matching. (I) Sib pairs, one-to-one matching. (J) Sib pairs, one-to-many matching with a query SNP profile. (K) Sib pairs, one-to-many matching with a query STR profile. (L) Sib pairs, needle-in-haystack matching. At each value of *c*, 100 fragmented genomic datasets are considered (Section 2.5). All rely on the same partition of the 1000 Genomes dataset into a test set with 75% of the individuals and a target set with the other 25% (Section 2.4); this partition corresponds to the median record-matching accuracy under one-to-one matching with the correctly specified hypothesis (Table 3). The pink line indicates the median of 100 trials with different fragmented datasets; the dashed lines around the pink line specify the minimum and maximum. The green and blue horizontal lines indicate the median record-matching accuracy using the full-coverage 1000 Genomes (Table 3) and HGDP datasets (Table S1), respectively.

For all four matching schemes, the median accuracy for 1000 Genomes exceeds the median accuracy for HGDP; numerical values for HGDP appear in Table S1 and for 1000 Genomes in Tables 2 and S2. As the coverage of 1000 Genomes decreases in fragmentary datasets starting from *c* = 0.9, accuracy decreases as well.

For one-to-one matching, decreasing the 1000 Genomes coverage *c* from 0.9, the median accuracy across 100 fragmentary SNP replicates begins at 1 at *c* = 0.9, remaining equal to 1 until the coverage reaches *c* = 0.06, for which the accuracy drops to 0.997 (Figure 3A). The HGDP median one-to-one accuracy of 0.991 (blue horizontal line) is achieved in 1000 Genomes at *c ≈* 0.05. The accuracy drops quickly after *c* = 0.03, with median accuracy 0.906; it is 0.677 at *c* = 0.02 and 0.181 at *c* = 0.01.

For one-to-many matching with a SNP query (Figure 3B) or STR query (Figure 3C), median accuracy drops somewhat faster than for one-to-one matching. Near ∼50% coverage (*c* = 0.5), it drops below 1, though it remains high at much lower coverage. The HGDP median accuracies (0.922, 0.940) are achieved at *c* ≈ 0.05.

For the needle-in-haystack scheme (Figure 3D), the median accuracy is still lower. The value drops below 0.9 at *c ≈* 0.3. The HGDP median accuracy for this scheme (0.532) is achieved at *c* ≈ 0.05.

### 3.2 Parent–offspring

When **Δ**_true_ **= Δ**_test_ = parent–offspring (Figure 3E-H), accuracies are lower than for the case in which SNP and STR profiles represent the same individual (Figure 3A-D). For all coverage levels *c* < 1, the median accuracy of one-to-one matching (pink line in Figure 3E) is lower than the match accuracy with the full dataset (0.738; green horizontal line). At *c* = 0.9, median accuracy is 0.725; it decreases to achieve the HGDP median one-to-one accuracy of 0.367 (blue horizontal line) at *c* ≈ 0.09. At *c* = 0.01, median accuracy is 0.032.

For one-to-many matching with a SNP query (Figure 3F) and one-to-many matching with an STR query (Figure 3G), the median accuracy drops faster from 0.649 and 0.649 at full coverage to 0.374 and 0.371 at *c* = 0.1, respectively. The HGDP median values (0.367, 0.358) are achieved at *c* ≈ 0.1.

For the needle-in-haystack scheme (Figure 3H), the median accuracy is below the full-coverage median accuracy for all coverage values, dropping from 0.113 at *c* = 0.9 to 0.026 at *c* = 0.1. The HGDP median needle-in-haystack accuracy (0.037) is achieved at *c ≈* 0.15.

### 3.3 Sib pairs

When **Δ**_true_ **= Δ**_test_ = sib pairs (Figure 3I-L), accuracies are comparable to those seen when the SNP and STR profiles represent a parent and offspring (Figure 3E-H) and lower than those in which they represent the same individual (Figure 3A-D). For all values of the coverage *c*, the median accuracy of one-to-one matching (Figure 3I) is lower than the accuracy from the full-coverage data (0.693; green horizontal line). At *c* = 0.9, the one-to-one median accuracy is 0.674, decreasing to achieve the HGDP median one-to-one accuracy (0.404; blue horizontal line) at *c* ≈ 0.11. At *c* = 0.01, the median accuracy is 0.038.

For one-to-many matching with a SNP query (Figure 3J) or STR query (Figure 3K), the median accuracy drops from a lower starting point, decreasing from 0.626 and 0.636 at full coverage to 0.383 and 0.383 at *c* = 0.1. The HGDP median one-to-many accuracies (0.394, 0.394) are achieved at *c* ≈ 0.11.

For the needle-in-haystack scheme (Figure 3L), the median accuracy lies below the full-coverage median accuracy for all coverage values we simulated, declining from 0.182 at *c* = 0.9 to 0.058 at *c* = 0.1. The HGDP median for the needle-in-haystack scheme (0.046) is achieved at *c* ≈ 0.08.

### 3.4 Ratio of posterior and prior odds

Using Eq. 4 in Section 2.2.2, Figure 4A and Table 4 display the minimum match score *λ* required to achieve a desired ratio of the posterior odds to the prior odds. For example, to obtain a posterior odds of a match equal to 10^4^ with prior odds 10^−9^, the match score under a specified test hypothesis (Eq. 1) must reach the threshold for a ratio of posterior odds to prior odds equal to 10^13^, or 29.93.

**Table 4:** The fraction of true matches with match score exceeding the minimum threshold for achieving a desired ratio of posterior and prior odds. The values correspond to those plotted in Figure 4.

| Ratio of posterior odds and prior odds |  |  |  |  |  |  |  |  |  |  |  |  |  |  |  |  |  |  |
| --- | --- | --- | --- | --- | --- | --- | --- | --- | --- | --- | --- | --- | --- | --- | --- | --- | --- | --- |
|  | 10<sup>0</sup> | 10<sup>1</sup> | 10<sup>2</sup> | 10<sup>3</sup> | 10<sup>4</sup> | 10<sup>5</sup> | 10<sup>6</sup> | 10<sup>7</sup> | 10<sup>8</sup> | 10<sup>9</sup> | 10<sup>10</sup> | 10<sup>11</sup> | 10<sup>12</sup> | 10<sup>13</sup> | 10<sup>14</sup> | 10<sup>15</sup> | 10<sup>16</sup> | 10<sup>17</sup> |
| Minimum match score |  |  |  |  |  |  |  |  |  |  |  |  |  |  |  |  |  |  |
|  | 0 | 2.30 | 4.61 | 6.91 | 9.21 | 11.51 | 13.82 | 16.12 | 18.42 | 20.72 | 23.03 | 25.33 | 27.63 | 29.93 | 32.24 | 34.54 | 36.84 | 39.14 |
| Fraction of true matches exceeding the minimum match score |  |  |  |  |  |  |  |  |  |  |  |  |  |  |  |  |  |  |
| Same individual | 0.99 | 0.99 | 0.98 | 0.96 | 0.95 | 0.93 | 0.89 | 0.85 | 0.81 | 0.75 | 0.67 | 0.58 | 0.47 | 0.37 | 0.27 | 0.19 | 0.12 | 0.09 |
| Parent-offspring | 0.88 | 0.74 | 0.58 | 0.35 | 0.18 | 0.05 | 0.02 | 0.00 | 0.00 | 0.00 | 0.00 | 0.00 | 0.00 | 0.00 | 0.00 | 0.00 | 0.00 | 0.00 |
| Sib pairs | 0.90 | 0.74 | 0.53 | 0.33 | 0.18 | 0.07 | 0.03 | 0.01 | 0.00 | 0.00 | 0.00 | 0.00 | 0.00 | 0.00 | 0.00 | 0.00 | 0.00 | 0.00 |

**Figure 4:**
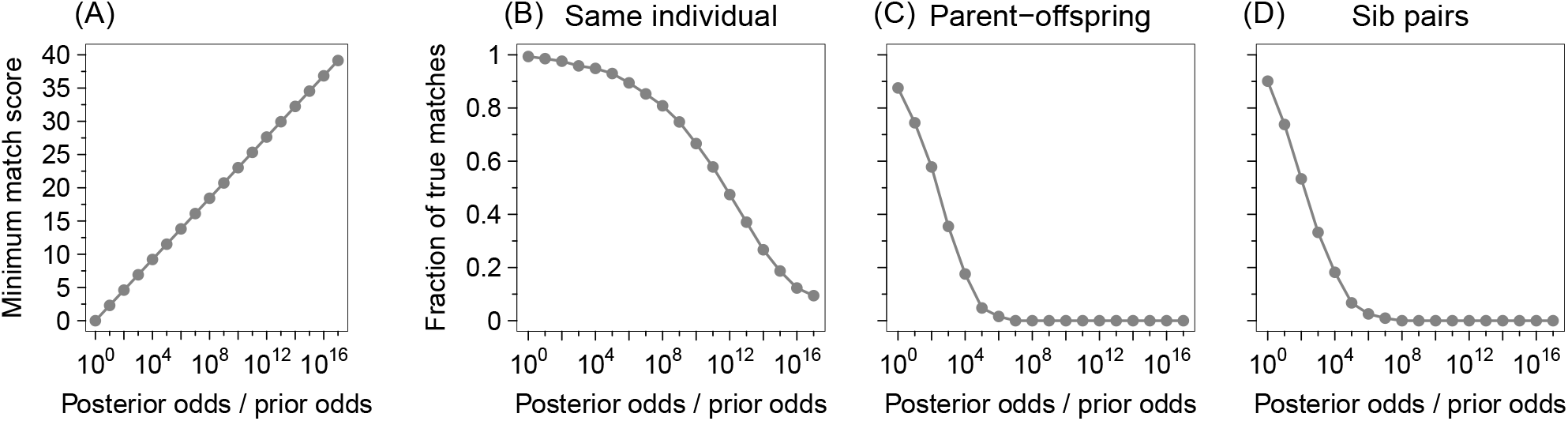
Numerical values of the fraction of true matches with match score exceeding the minimum threshold for achieving a desired ratio of posterior and prior odds. (A) The minimum match score required to achieve a desired ratio of posterior and prior odds (Eq. 4). (B)–(D) The fraction of true matches with match score exceeding the minimum match score required for achieving a desired ratio of posterior and prior odds (shown in panel A) when **Δ**_true_ = **Δ**_test_ and *c* = 1. The values (Table 4) were computed from 626 individuals in the test set of the partition corresponding to the median one-to-one match accuracy under full genomic SNP coverage using the 1000 Genomes dataset (Section 2.4 and Table 3).

For each relatedness scenario and a ratio of posterior and prior odds, we computed the fraction of true matches whose match score exceeded the minimum match score (Figure 4A) required for achieving a prescribed ratio of posterior to prior odds. When **Δ**_true_ = **Δ**_test_ = same individual (Figure 4B), if the prior probability of two individuals being unrelated is 10^10^ times that of them being the same individual (prior odds 10^*−*10^), then 67% of true matches achieve posterior odds exceeding 1 (ratio 10^10^ for posterior and prior odds), and 9% achieve posterior odds greater than 10^7^ (ratio 10^17^). When the prior is uninformative, with prior odds of 1, 99% of true matches exceed the match score required for attaining posterior odds of 1 (ratio 10^0^), and 85% of true match pairs have posterior odds greater than 10^7^ (ratio 10^7^).

When **Δ**_true_ = **Δ**_test_ = parent–offspring (Figure 4C), the behavior is similar to the same-individual case, but with lower probabilities of attaining specified thresholds. A high value of 88% of true matches achieve posterior odds 1 with prior odds of 1 (ratio 10^0^). However, in contrast to the same-individual case, posterior odds values in the range of [10^0^, 10^11^] are largely unattainable with prior odds below 10^*−*6^ (ratios 10^6^ to 10^17^). Finally, when **Δ**_true_ = **Δ**_test_ = sib pairs (Figure 4D), patterns are similar to the parent–offspring case.

### 3.5 Misspecified hypotheses Δ_true_ ≠ Δ_test_

Figure S1 and Table S2 examine the record-matching accuracies for six pairs of misspecified relatedness hypotheses, **Δ**_true_ ≠ **Δ**_test_. Accuracies are generally smaller for the misspecified hypotheses than for the correctly specified hypotheses. When profiles represent the same individual (**Δ**_true_ = same individual), the misspecified parent–offspring (**Δ**_test_ = parent–offspring) and sib-pair hypotheses (**Δ**_test_ = sib pairs) continue to detect the relationship with relatively high accuracy. Misspecifying a parent–offspring pair (**Δ**_true_ = parent– offspring) as sibs (**Δ**_test_ = sib pairs) or a sib pair (**Δ**_true_ = sib pairs) as parent–offspring (**Δ**_test_ = same individual) has a smaller effect on record-matching accuracy than misspecifying either type of pair (**Δ**_true_ = parent–offspring, **Δ**_true_ = sib pairs) as arising from the same individual (**Δ**_test_ = same individual).

## 4 Discussion

We have applied genetic record-matching to fragmentary SNP data, comparing the resulting record-matching accuracies to those obtained with a genome-sequence dataset and to those seen in our previous analysis of data with lower marker density. We have found that for the 1000 Genomes data, representing fully sequenced genomes, record-matching accuracies exceed those seen previously in the Human Genome Diversity Panel, typed with SNP genotyping arrays (Figure 3). Hence, accuracies at the level seen previously with genotyping arrays can be obtained in genome sequencing with incomplete coverage, often at a level of 5-10% of the genome (Figure 3). When matching profiles from the same individual, record-matching accuracy with the full genome is high in all four matching schemes tested—one-to-one, one-to-many with a SNP query, one-to-many with an STR query, and needle-in-haystack. When SNP and STR profiles are matched via one-to-one matching, the record-matching accuracy seen with the full genome is obtained with genomic overage as low as 6%.

The results demonstrate that the accuracy of genetic record-matching can be increased substantially beyond what we previously reported [16, 17]. In a calculation designed to test if a true match could be detected in a population the size of the United States adult population at posterior probability 10*/*11, Edge et al. [16] found that with 17-locus profiles, 8% of SNP–STR profile pairs matched closely enough that the true match would be detected at a threshold of 2.3 × 10^9^ for the ratio of posterior to prior odds. In our larger dataset, we find that even with smaller 15-locus profiles, a larger 67% of pairs would be detected at a more stringent threshold of 10^10^ for the ratio of posterior to prior odds (Table 4). This result indicates a sizeable probability that a true match of interest could be identified by genetic record-matching with high confidence via a query of a SNP database with an STR profile, or vice versa, even in a large population.

The improvement in accuracy that we have observed arises from multiple factors. First, our 1000 Genomes analysis makes use of sequenced genomes, so that the SNP density is substantially increased beyond the earlier HGDP SNP-genotyping studies [16, 17]. Second, the 1000 Genomes dataset contains more individuals than the HGDP dataset, so that the increased sample size might also have contributed to the increased accuracy. Indeed, in an additional analysis in Figure 5, which examines record-matching accuracy in subsamples of the 1000 Genomes dataset with varying sample size, considering full genomic coverage (*c* = 1) and searching for same-individual matches, we can observe that particularly for the needle-in-haystack scheme, record-matching accuracy increases with increasing sample size. This pattern suggests that increasing the number of individuals in the reference panel to improve the estimation of genotype probabilities in Eq. 3 (“improving the needle detector”) may have a large enough effect in increasing record-matching accuracy to counteract the increase in the number of possible pairs among which matches are sought (“enlarging the haystack”).

**Figure 5:**
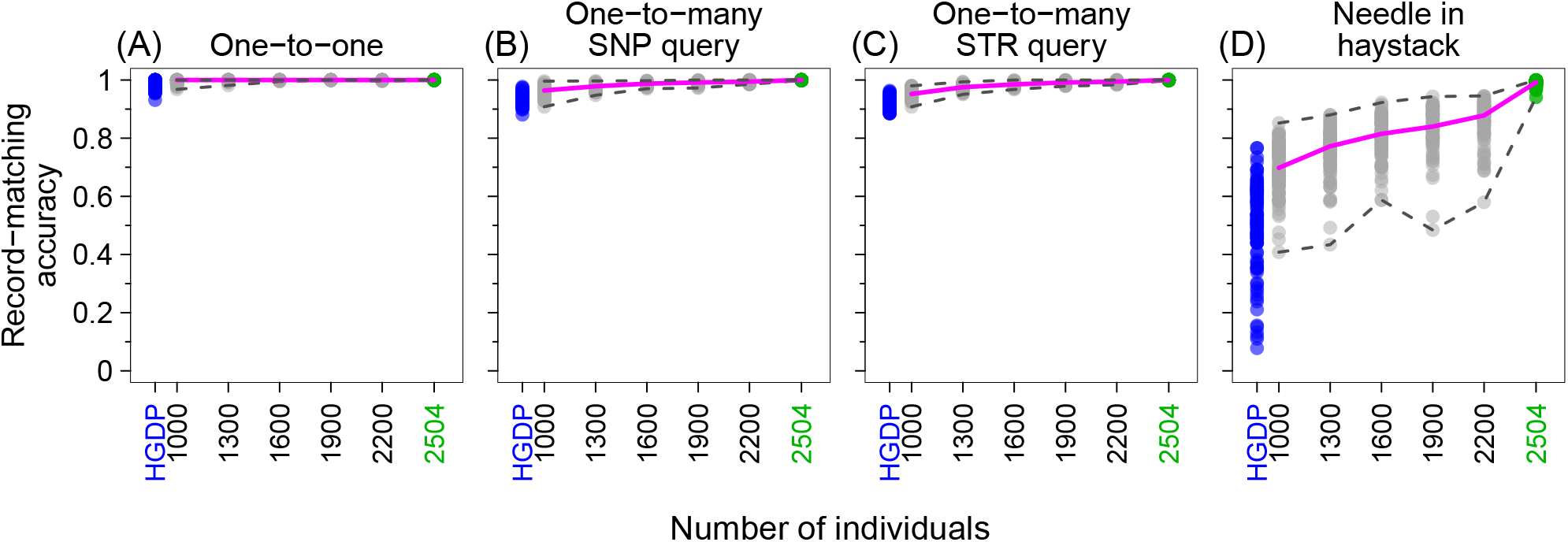
Record-matching accuracy in subsamples of varying size. The figure uses sampled individuals from the 1000 Genomes with full coverage (*c* = 1), considering 15 Codis loci and **Δ**_true_ **= Δ**_test_ = same individual. For each number of individuals in {1000, 1300, 1600, 1900, 2200}, we randomly sampled 100 sets of individuals from 2,504 individuals in the 1000 Genomes dataset and performed record matching on the reduced dataset, choosing 75% of the individuals for the training set and 25% for the test set. Green points consider all 2,504 individuals in the 1000 Genomes and show the values for the 100 replicates summarized in Table 3. The pink line indicates the median one-to-one matching accuracy of 100 trials. For comparison, the blue points indicate the corresponding results using the full HGDP dataset of 872 individuals, reporting the values for the 100 replicates summarized in the upper left corner of Table S1.

We note that our analysis did not distinguish profiles by source population, considering the entire reference panel as a single population. Conducting record-matching separately in different populations by relying on relevant reference panels for imputation in different subgroups [25–27] could potentially increase the imputation accuracy of STRs from SNPs—and by consequence, the record-matching accuracy. In cases in which population membership of a profile of interest is unknown, such a procedure could identify suitable imputation reference panels via genetic ancestry inference for a query profile prior to applying record-matching [28].

We also note that to facilitate comparisons between the 1000 Genomes and HGDP, our analyses relied on the 15 Codis loci present in both datasets. As in analyses of Edge et al. [16] and Kim et al. [17] that examined increasingly large numbers of loci, however, we expect that the record-matching accuracy would improve for a given genomic coverage if it were possible to include all 20 Codis loci.

A third limitation is that we used a simple simulation to produce fragmentary genomic SNP datasets, assuming that for a given coverage *c*, SNPs amplify independently of one another, and that amplification patterns are independent across individuals. With actual degraded DNA, fragmentary datasets are likely to possess spatial correlation across the genome, containing multiple neighboring SNPs genotyped on the same fragments of DNA (Figure 1B). Inclusion of spatial correlation would increase the probability that at a given coverage, in some individuals some STRs would possess no neighboring genotyped SNPs, so that no information would be available to impute those STR genotypes. Hence, high levels of spatial correlation in amplification for a fixed coverage could reduce record-matching accuracy compared to the model we have considered, especially at coverage levels low enough to eliminate all SNPs around some of the STRs. The simple simulation approach we have considered is intended to be informative about the general setting of fragmentary genomic SNP data; as degradation and amplification patterns are likely to differ for different DNA sources, details of the simulation could be considered specifically based on such patterns.

The results have a clear application in settings in which an investigator would have genotyped STRs and tested those STRs for matches against an STR database, were STRs possible to genotype. If samples are degraded to the point where only SNPs and not STRs can be obtained—as might occur in settings of older criminal-justice samples, mass disasters, burned material, or ancient DNA—then our approach could be used to test the resulting SNP profile against an STR database. In such cases, the genetic record-matching technique is used simply to overcome the technical challenge of genotyping STRs in degraded material—in existing investigative settings, rather than by introducing new investigative settings.

As discussed by Edge et al. [16] and Kim et al. [17], however, genetic record-matching can also produce new types of information linkages if investigators or others possess access to both SNP and STR databases. Profiles in different databases could in principle be connected if participants in a biomedical or genealogical SNP database or their close relatives appear in a forensic STR database. Our results increase the potential accuracy for such efforts. Thus, the study contributes to a growing body of work on cross-database linkages involving genetic data, with significant investigative potential and with attendant privacy risks [29, 30]. Our previous work [16, 17] discussed privacy risks from the linkages between genetic databases—and possibly even phenotype databases—that are enabled by genetic record matching, and even before the 2018 introduction of the long-range search technique combining genetic and genealogical data, Edge et al. [16] wrote *“Contrary to the view that* Codis *genotypes expose no phenotypes, a* Codis *profile on a person together with a SNP database—if the person is in the database—in principle may contain all of the phenotypic information that can be reliably predicted from the SNP record. Conversely, participants in biomedical research or personal genomics who have consented to share their SNP genotypes may be subject to a previously unappreciated risk: identification in a forensic STR database*.*”*

The increased record-matching accuracy that we have detected in a larger, denser dataset than that used by Edge et al. [16] and Kim et al. [17] only magnifies the privacy concern. The potential for employing genetic record-matching to use one type of individual-level information to reveal information of another type enhances both the potential uses of the technique for individual identification in degraded crime-scene samples, ancient samples, missing-persons and mass-disasters cases—as well as the potential risks that excess information could be revealed, either by an authorized user or by an attacker. Further consideration is needed of the benefits and privacy risks emerging from cross-database linkages involving SNPs and STRs—and phenotypes.

## Acknowledgments

We acknowledge support from National Institutes of Health grant R01 HG005855.

---

**Figure S1:**
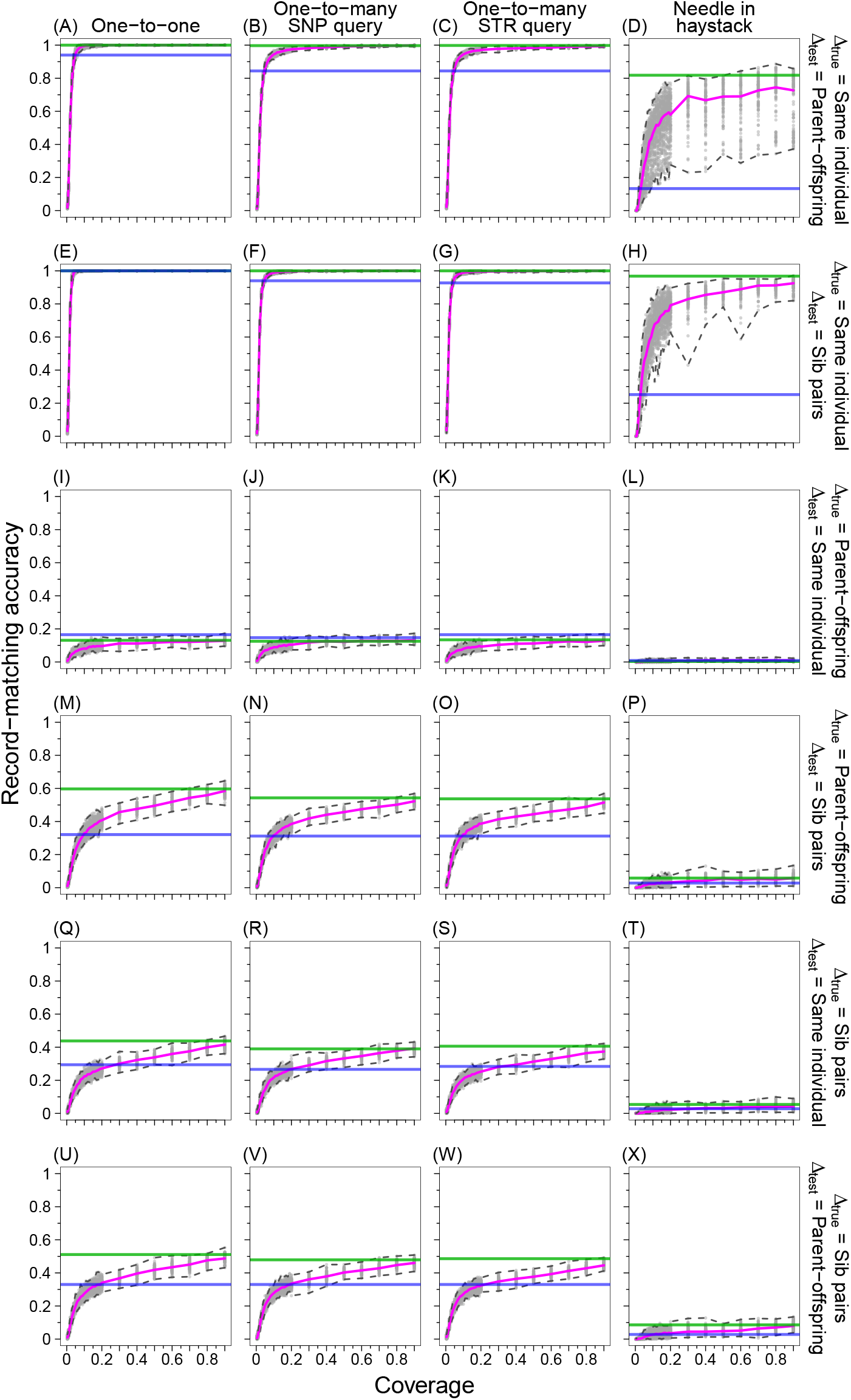
Record-matching accuracy in fragmented genomic data as a fraction of the genomic coverage *c*, for Δ_true_ ≠ Δ_test_. (A-D): **Δ**_true_ = same individual, **Δ**_test_ = parent–offspring. (E-H): **Δ**_true_ = same individual, **Δ**_test_ = sib pairs. (I-L): **Δ**_true_ = parent–offspring, **Δ**_test_ = same individual. (M-P): **Δ**_true_ = parent–offspring, **Δ**_test_ = sib pairs. (Q-T): **Δ**_true_ = sib pairs, **Δ**_test_ = same individual. (U-X): **Δ**_true_ = sib pairs, **Δ**_test_ = parent–offspring. The figure design follows Figure 3.

**Table S1:** Record-matching accuracies using the HGDP dataset and 15 CODIS loci. The table summarizes 100 partitions into training and test sets, applying record matching to the HGDP dataset with no missing data. The STRs used are listed in Table 2.

| $\Delta_{\text{true}} \backslash \Delta_{\text{test}}$ | Same individual | | Parent-offspring | | Sib pairs | | Match-assignment scheme |
| --- | --- | --- | --- | --- | --- | --- | --- |
|  | Median | Min, Max | Median | Min, Max | Median | Min, Max |  |
| Same individual | 0.991 | 0.945, 1.000 | 0.940 | 0.858, 0.991 | 1.000 | 0.963, 1.000 | One-to-one |
|  | 0.922 | 0.885, 0.972 | 0.844 | 0.789, 0.899 | 0.940 | 0.881, 0.972 | One-to-many: SNP query |
|  | 0.940 | 0.899, 0.982 | 0.844 | 0.775, 0.894 | 0.927 | 0.881, 0.972 | One-to-many: STR query |
|  | 0.532 | 0.055, 0.803 | 0.133 | 0.005, 0.358 | 0.252 | 0.046, 0.596 | Needle-in-haystack |
| Parent-offspring | 0.165 | 0.064, 0.248 | 0.367 | 0.266, 0.495 | 0.321 | 0.220, 0.422 | One-to-one |
|  | 0.147 | 0.083, 0.229 | 0.367 | 0.220, 0.459 | 0.312 | 0.211, 0.404 | One-to-many: SNP query |
|  | 0.165 | 0.092, 0.248 | 0.358 | 0.257, 0.450 | 0.312 | 0.202, 0.431 | One-to-many: STR query |
|  | 0.009 | 0.000, 0.037 | 0.037 | 0.000, 0.138 | 0.028 | 0.000, 0.128 | Needle-in-haystack |
| Sib pairs | 0.294 | 0.156, 0.440 | 0.330 | 0.229, 0.450 | 0.404 | 0.284, 0.550 | One-to-one |
|  | 0.266 | 0.174, 0.358 | 0.330 | 0.211, 0.431 | 0.394 | 0.294, 0.505 | One-to-many: SNP query |
|  | 0.284 | 0.193, 0.385 | 0.330 | 0.202, 0.459 | 0.394 | 0.275, 0.495 | One-to-many: STR query |
|  | 0.028 | 0.000, 0.083 | 0.028 | 0.000, 0.119 | 0.046 | 0.009, 0.147 | Needle-in-haystack |

**Table S2:** Record-matching accuracies using the 1000 Genomes dataset and 15 CODIS loci. The table summarizes 100 partitions into training and test sets, applying record matching to the 1000 Genomes dataset with no missing data. The STRs used are listed in Table 2. The block diagonal entries with **Δ**_true_ **= Δ**_test_ also appear in Table 3.

| $\Delta_{\text{true}} \backslash \Delta_{\text{test}}$ | Same individual | | Parent-offspring | | Sib pairs | | Match-assignment scheme |
| --- | --- | --- | --- | --- | --- | --- | --- |
|  | Median | Min, Max | Median | Min, Max | Median | Min, Max |  |
| Same individual | 1.000 | 1.000, 1.000 | 1.000 | 1.000, 1.000 | 1.000 | 1.000, 1.000 | One-to-one |
|  | 1.000 | 0.998, 1.000 | 0.997 | 0.990, 1.000 | 1.000 | 0.998, 1.000 | One-to-many: SNP query |
|  | 1.000 | 0.998, 1.000 | 0.998 | 0.994, 1.000 | 1.000 | 0.998, 1.000 | One-to-many: STR query |
|  | 0.992 | 0.941, 1.000 | 0.818 | 0.492, 0.922 | 0.968 | 0.898, 0.995 | Needle-in-haystack |
| Parent-offspring | 0.131 | 0.083, 0.204 | 0.738 | 0.681, 0.812 | 0.597 | 0.514, 0.668 | One-to-one |
|  | 0.125 | 0.080, 0.169 | 0.649 | 0.581, 0.700 | 0.543 | 0.457, 0.613 | One-to-many: SNP query |
|  | 0.134 | 0.099, 0.192 | 0.649 | 0.597, 0.719 | 0.537 | 0.470, 0.610 | One-to-many: STR query |
|  | 0.003 | 0.000, 0.032 | 0.112 | 0.016, 0.256 | 0.058 | 0.003, 0.150 | Needle-in-haystack |
| Sib pairs | 0.438 | 0.361, 0.527 | 0.511 | 0.447, 0.585 | 0.693 | 0.623, 0.773 | One-to-one |
|  | 0.390 | 0.335, 0.470 | 0.479 | 0.412, 0.527 | 0.626 | 0.546, 0.674 | One-to-many: SNP query |
|  | 0.406 | 0.351, 0.476 | 0.486 | 0.425, 0.546 | 0.636 | 0.572, 0.703 | One-to-many: STR query |
|  | 0.054 | 0.010, 0.115 | 0.086 | 0.006, 0.169 | 0.160 | 0.019, 0.275 | Needle-in-haystack |

